# Elucidating Variants in Systemic JIA from Analysis of Illumina & Ion Torrent Exome Data

**DOI:** 10.1101/023580

**Authors:** M. Rahman

## Abstract

**Motivation:** Children with systemic juvenile idiopathic arthritis (SJIA) are affected by a wide-range of complications. Partially arising from the difficulty of diagnosis due to the idiopathic nature of the indication. There may be a genetic basis for SJIA, which could help in both diagnosis, and treatment.

**Results:** Two mutations in the Fc epsilon RI pathway, including PIK3CD, were detected in low-coverage Ion Torrent data. Variants of unknown significance were detected within HLA regions on standard Illumina exomes. CSF2RA, which could account for pulmonary observations, had insignificant coverage on both datasets.

**Availability:** ftp.systemicjia.com

## 1 INTRODUCTION

SJIA is an autoimmune disease with additional extra-articular symptoms including pulmonary complications. Because the exact cause of the disease remains a mystery, exome sequencing was applied in this investigation. Data was generated for a sample-set of four individuals, including one child affected by SJIA, a sibling, and both biological parents.

## 2 APPROACH

A hybrid assembly combining Ion Torrent and Illumina, unenriched short reads was used to generate the low coverage datasets. Meanwhile, a standard Illumina exome using TrueSeq enrichment was generated by vendor Ambry Genetics. Both datasets contain the same pedigree and were analyzed with a custom NGS pipeline, as well as the GATK best practices (Van der Auwera *et al*., 2013).

## 3 METHODS

Samples were collected from saliva or buccal swabs, using Oragene DNA OG-575 or OG-500, by DNA Genotek. Libraries were prepared using the Wafergen Apollo 324 system. Sequencing performed on Ion 318 chips with kit v2, and Illumina HiSeq2500. Generated FASTQ files processed with FASTX-Toolkit. Reads aligned with BWA v0.7.1(Li H., 2013), using reference UCSC hg19; BAM files were further processed by Picard Tools.

VCF files were generated via two methods, Samtools mpileup and GATK HaplotypeCaller. Reference SNP IDs (RSID) were added from dbSNP build 138. Functional annotations performed with two seperate algorithms, SnpEff v4.1 and Ensembl Variant Effect Predictor (VEP) with the Loss of Function Transcript Effect Estimator (LOFTEE) plugin. Pathway and disease annotations were added from KEGG, ClinVar, and OMIM.

Variants above a certain quality threshold were compared within the pedigree. Only those data points which are unique for individuals affected with SJIA, but not present in a homozygous manner in the rest of the pedigree, move to the next step of analysis. If a variant is predicted to change the structure of a protein, its role in a molecular pathway is determined. Finally, what disease would result if such a break in a given pathway were to occur is deduced. Currently, the component of the pipeline performing these tasks is being called D^3^ (Definitive Desired Diagnosis).

## 4 DISCUSSION

Immunoglobulin E (IgE) plays an important role in allergy disorders and parasite immunity, and the Fc region is particularly key to mast cells. Table 1 illustrates the two mutations found to be possibly affecting these functions. While the standard Illumina exome shows PIK3CD to be heterozygous with depth of read of 14, and short reads of less than 100 bp; hybrid Ion Torrent data shows homozygous mutation with 6/6 reads with lengths between 150-200 bp.

**Table 1.** Fc*ϵ*RI (high-affinity IgE receptor) Mutations

| Chr | Position | RSID | Ref/Alt | Gene | Effect |
| --- | --- | --- | --- | --- | --- |
| 1 | 9777599 | rs61755420 | C/G | PIK3CD | missense |
| 22 | 31532960 | rs2232183 | C/T | PLA2G3 | missense |

TRANSCRIPT\_ID=NM\_005026.3 missense SER/CYS
TRANSCRIPT\_ID=NM\_015715.4 missense ARG/GLN

Human leukocyte antigen (HLA) system consists of a set of genes which regulate the immune system. These regions are also known to be highly variable between individuals. In the Ion Torrent hybrid data HLA regions were covered without bias, and variants for affected individuals were similar to those of the pedigree. Standard Illumina exome data focused more on HLA regions, and variants unique to affected individual, including in comparison to twin, were prevalent. Although their significance remains unknown.

**Fig. 1.**
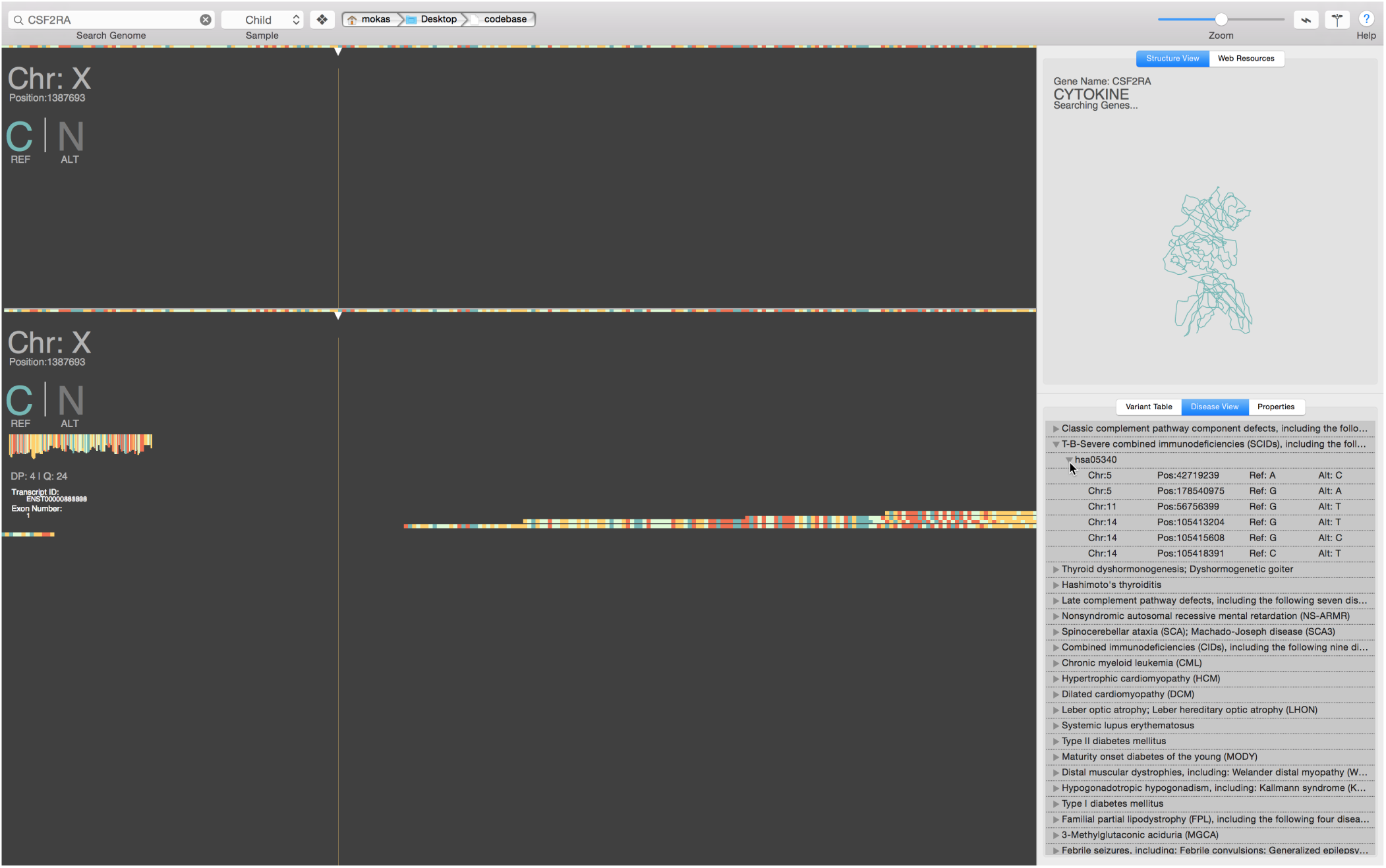
CSF2RA gene in screenshot from viewer showing both datasets. Standard Illumina exome (top) shows no reads, as the gene was likely not included in the kit library. Hybrid Ion Torrent, low coverage (bottom) shows some reads in the region, though large gaps are still prevalent.

## 5 CONCLUSION

Sequence alignments created here are from two different platforms, as well as being enriched and unenriched. They cover a wide range of the genome and have provided some direction. Low coverage Ion Torrent hybrid data shows variants of interest in the Fc*ϵ*RI pathway, which requires additional data to be confident. Enriched Illumina exome shows many variants in the HLA regions, though their significance is unknown. Both datasets leave huge gaps in the genome as well as inadequate reads in other regions to make confident variant calling.

Observed pulmonary complications and biopsy results, made granulocyte macrophage colony-stimulating factor receptor (CSF2RA) worth investigation. Standard Illumina exome data completely missed this gene, while the hybrid Ion Torrent data did not detect any pathogenic variants with the sparse reads in the dataset.

While the hybrid Ion Torrent data provided longer reads, nearly double of Illumina alone, the depth of read was low; resulting in larger overlaps and longer contigs, but lower quality base calls. Reads mapped in an expected manner for the standard Illumina exome, which is well characterized. Regions predetermined for the TrueSeq enrichment kit provided good coverage, but were less helpful due to the idopathic nature of SJIA.

Additional data will help to elucidate the complex nature of this immunological disorder. Whether a genomic cause exists in single genes or pathways, or more difficult to interpret regions like HLA, the current datasets provide a good starting point.

## ACKNOWLEDGEMENT

This work used the Vincent J. Coates Genomics Sequencing Laboratory at UC Berkeley, supported by NIH S10 Instrumentation Grants S10RR029668 and S10RR027303.

## REFERENCES

Van der Auwera GA, Carneiro M, Hartl C, Poplin R, del Angel G, Levy-Moonshine A, Jordan T, Shakir K, Roazen D, Thibault J, Banks E, Garimella K, Altshuler D, Gabriel S, DePristo M (2013) From FastQ Data to High-Confidence Variant Calls: The Genome Analysis Toolkit Best Practices Pipeline Current Protocols In Bioinformatics, 43, 11.10.1–11.10.33.

Li H. (2013) Aligning sequence reads, clone sequences and assembly contigs with BWA-MEM, arXiv:1303.3997v2, [q-bio.GN]

Cingolani, P., Platts, A., Wang, L. L., Coon, M., Nguyen, T., Wang, L., Ruden, D. M. (2012) A program for annotating and predicting the effects of single nucleotide polymorphisms, SnpEff: SNPs in the genome of Drosophila melanogaster strain w1118; iso-2; iso-3, Fly, 2, 80–92.

Kanehisa, M., Goto, S., Sato, Y., Kawashima, M., Furumichi, M., and Tanabe, M. (2014) Data, information, knowledge and principle: back to metabolism in KEGG, Nucleic Acids Res., 42, D199–D205.

